# Diflunisal targets the HMGB1/CXCL12 heterocomplex and blocks immune cell recruitment

**DOI:** 10.1101/563890

**Authors:** Federica De Leo, Giacomo Quilici, Mario Tirone, Valeria Mannella, Francesco De Marchis, Alessandro Preti, Alessandro Gori, Maura Casalgrandi, Rosanna Mezzapelle, Marco E. Bianchi, Giovanna Musco

## Abstract

Extracellular HMGB1 triggers inflammation following infection or injury, and supports tumorigenesis in inflammation-related malignancies. HMGB1 has several redox states: reduced HMGB1 recruits inflammatory cells to injured tissues forming a heterocomplex with CXCL12 and signaling *via* its receptor CXCR4; disulfide-containing HMGB1 binds to TLR4 and promotes inflammatory responses. Here we show that Diflunisal, an aspirin-like nonsteroidal anti-inflammatory drug (NSAID) that has been in clinical use for decades, specifically inhibits *in vitro* and *in vivo* the chemotactic activity of HMGB1 at nanomolar concentrations, at least in part by binding directly to both HMGB1 and CXCL12 and disrupting their heterocomplex. Importantly, Diflunisal does not inhibit TLR4-dependent responses. Our findings clarify the mode of action of Diflunisal, and open the way to the rational design of functionally specific anti-inflammatory drugs.

## Introduction

High-Mobility Group Box-1 (HMGB1), a highly conserved and abundant nuclear protein expressed in all eukaryotic cells, is a key trigger of inflammation (Andersson & Tracey, 2011; Bianchi *et al*, 2017). HMGB1 consists of two structurally independent L-shaped tandem HMG-box domains (∼80 amino acids each), referred to as Box A and Box B, connected by a flexible linker and followed by a long acidic C-terminal tail of 30 amino acids (Thomas, 2001). Within the nucleus, HMGB1 is a DNA chaperone that regulates nucleosome assembly and chromatin structure, promotes interactions between several transcription factors and facilitates their binding to DNA (Štros, 2010). HMGB1 is passively released in the extracellular space by dead cells and is actively secreted by severely stressed cells (reviewed in Bianchi *et al*, 2017). Extracellular HMGB1 alerts the host to unscheduled cell death, to stress, and to microbial invasion, thus playing a key role in inflammation and immune responses (Bianchi *et al*, 2017). In particular, it triggers inflammation following injury or infection by first recruiting inflammatory cells (Schiraldi *et al*, 2012) and then activating them (Venereau *et al*, 2012). HMGB1-induced recruitment of inflammatory cells depends on the formation of a heterocomplex with the chemokine CXCL12, which in turn activates CXCR4, a G-protein coupled receptor (Venereau *et al*, 2012). Notably, the formation of a disulfide bond between cysteines 22 and 44 makes HMGB1 unable to form the heterocomplex, but makes it a ligand of TLR4, a receptor that also recognizes LPS (Venereau *et al*, 2012). Since the two redox states of HMGB1 have alternative activities, it would be desirable to inhibit them selectively.

Interfering with the formation of the HMGB1-CXCL12 heterocomplex with small molecules is an attractive but challenging route, mainly because the large and flat interfaces typically associated with protein-protein interactions have reduced druggability (Laraia *et al*, 2015; Scott *et al*, 2016). Here, we show that Diflunisal (DFL) (Fig. 1A), an aspirin-like nonsteroidal anti-inflammatory drug that was identified in the 1970s through phenotypic screens (Steelman *et al*, 1978), exerts its anti-inflammatory action at least in part by disrupting the interaction between HMGB1 and CXCL12.

**Figure 1.**
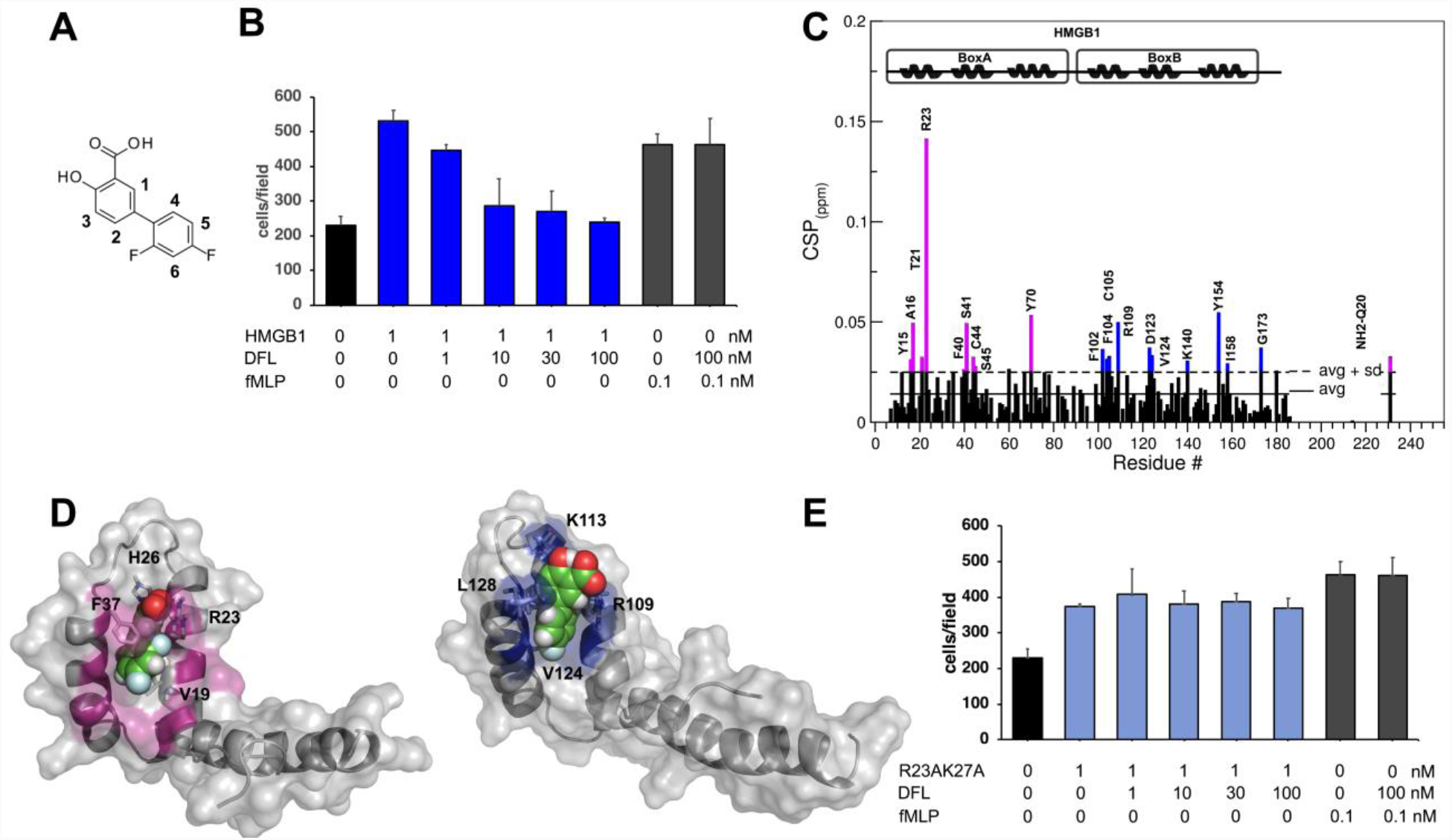
A) Chemical structure of DFL. B) DFL inhibits HMGB1-induced, but not fMLP-induced cell migration. 3T3 fibroblasts were subjected to chemotaxis assays in Boyden chambers, 1 nM HMGB1 or no chemoattractant was added in the lower chamber, together with the indicated concentrations of DFL. Data represent the average ± standard deviation (avg ± sd, n=3) of one representative experiment. C) Histogram showing residue-specific CSPs of ^15^N-labelled HMGB1 (∼0.1 mM) upon addition of ten-fold excess of DFL (helices are schematically represented on the top). Missing residues are prolines or are absent because of exchange with the solvent. Box A and Box B residues with CSP > avg + sd are represented in magenta and blue, respectively. D) HADDOCK models of interaction of DFL (CPK representation) with Box A (left) and Box B (right) (color, residues with CSP > avg + sd). HMGB1 residues (sticks) involved in hydrophobic and electrostatic interactions with DFL are explicitly labelled. E) DFL does not affect cell migration induced by fMLP nor by 1 nM R23AK27A HMGB1 mutant (compare to panel B). The data represent the avg ± sd of three replicates of one representative experiment. The difference in migration towards R23AK27A HMGB1 in the presence or absence of DFL is statistically non-significant (one-way ANOVA plus post tests).

## Results and Discussion

We recently found that Salicylic Acid suppresses HMGB1’s pro-inflammatory activity and inhibits HMGB1-induced motility, migration, invasion and anchorage-independent colony formation of mesothelioma cells *via* a cyclooxygenase-2–independent mechanism (Choi *et al*, 2015; Yang *et al*, 2015). Aiming at finding more potent HMGB1 inhibitors, we tested DFL, another FDA-approved salicylate, using migration of mouse 3T3 fibroblasts as cellular readout (Fig. 1B). DFL halved the number of migrating cells already at a concentration of 10 nM, the lowest documented IC_50_ so far for HMGB1 inhibitors (Venereau *et al*, 2016). Importantly, DFL did not affect chemotaxis toward fMLP (Fig. 1B) indicating that it does not influence the general mobility of fibroblasts.

We then asked whether DFL was a direct ligand of HMGB1. We performed NMR saturation transfer difference (STD) (Mayer & Meyer, 2001) and water-Ligand Observation via Gradient SpectroscopY (waterLOGSY) experiments (Dalvit *et al*, 2001) on DFL (1 mM) in the presence of sub-stoichiometric concentrations of recombinant HMGB1 (50 μM). We observed STD signals and inversion of the sign in waterLOGSY spectra for all DFL protons to varying extents, thus demonstrating a direct DFL-HMGB1 interaction (Appendix Fig. S1).

To identify the DFL binding site on HMGB1 we monitored perturbations of its ^1^H-^15^N Heteronuclear Single Quantum Coherence (HSQC) spectrum upon DFL addition. Binding occurred in fast-intermediate regime on the NMR time scale (Appendix Fig. S2). The pattern of chemical shift perturbations (CSPs) indicated that DFL binds to both boxes in full-length HMGB1 (Fig. 1C). Indeed, residues whose amide resonances showed significant CSPs (CSP > avg + sd), *i.e.* Y15, A16, Q20, T21, R23, F40, S41, C44, S45 in Box A, and F102, F104, C105, R109, D123, V124, K140 in Box B, were similar to the ones mostly affected in DFL titrations into isolated HMG Boxes (Appendix Fig. S3). When mapped on the structure of the two HMG boxes, residues with significant CSP defined a surface located at the crux of the typical L-shaped fold, characterized by a small solvent-exposed hydrophobic surface suitable for favorable van der Waals (vdW) interactions with the aromatic rings of DFL (Fig. 1D).

CSPs data were next used to generate HADDOCK (Dominguez *et al*, 2003) data-driven docking models of DFL in complex with Box A and Box B. The binding modes are highly reminiscent of the ones observed for Glycyrrhizin (Mollica *et al*, 2007), a well-known HMGB1 inhibitor. DFL accommodates at the junction of the two arms of both individual HMG boxes, establishing favorable vdW interactions with the hydrophobic side chains of V19 and F37 in Box A and of V124 and L128 in Box B (Fig. 1D). Additionally, the models predict stabilizing electrostatic interactions between DFL carboxylate and R23 and R109 sidechains (Fig. 1D). To prove the role of electrostatic interactions in ligand binding, we took advantage of an HMGB1 mutant (R23AK27A) that was previously used to prove the binding of Salicylic Acid to the same interaction surface (Choi *et al*, 2015). Indeed, NMR titration of DFL into ^15^N labelled HMGB1 R23AK27A mutant induced significantly lower CSPs as compared to the ones observed for the wildtype protein (Appendix Fig. S4), indicative of a weaker binding. Accordingly, DFL was not effective on cell migration induced by the R23AK27A HMGB1 mutant (Fig. 1E).

Microscale thermophoresis (MST) measurements (Wienken *et al*, 2010), performed by titrating DFL into fluorescently labelled HMGB1, yielded an apparent dissociation constant (K_d_) of 1.6 ± 0.8 mM (Appendix Fig. S5), in agreement with the fast-intermediate exchange regime observed in NMR titrations. However, such low affinity was in striking contrast with the high activity of DFL in cell migration experiments, thus arguing for a more complex inhibition mechanism than a conventional ligand-protein binary interaction. We thus hypothesized that DFL might target the HMGB1/CXCL12 heterocomplex, rather than HMGB1 alone. To explore this possibility, we first investigated whether DFL could be a direct ligand of CXCL12. NMR titration experiments of DFL into 0.1 mM ^15^N labelled CXCL12 clearly demonstrated an interaction between CXCL12 and DFL in fast/intermediate exchange regime, with an apparent K_d_ of 800 ± 102 µM (Appendix Fig. S6). The highest resonance shifts involved residues located on the β1 strand (V23, H25, K27) and residues (A40, N45, Q48, V49) around the so-called *sY21_CXCR4_* binding site (Veldkamp *et al*, 2006, 2008), which is recognized by the sulfated CXCR4 extracellular N-terminus (Fig. 2A). Prompted by this observation, we performed STD and waterLOGSY competition experiments between DFL and the CXCR4_1-38sY21_ peptide that corresponds to the N-terminal tail of CXCR4 sulfated in position Y21 (Veldkamp *et al*, 2006, 2008). Reduction of DFL signal intensities in STD and waterLOGSY spectra upon addition of CXCR4_1-38sY21_ indicated that both ligands compete for the same binding site (Fig. 2B).

**Figure 2.**
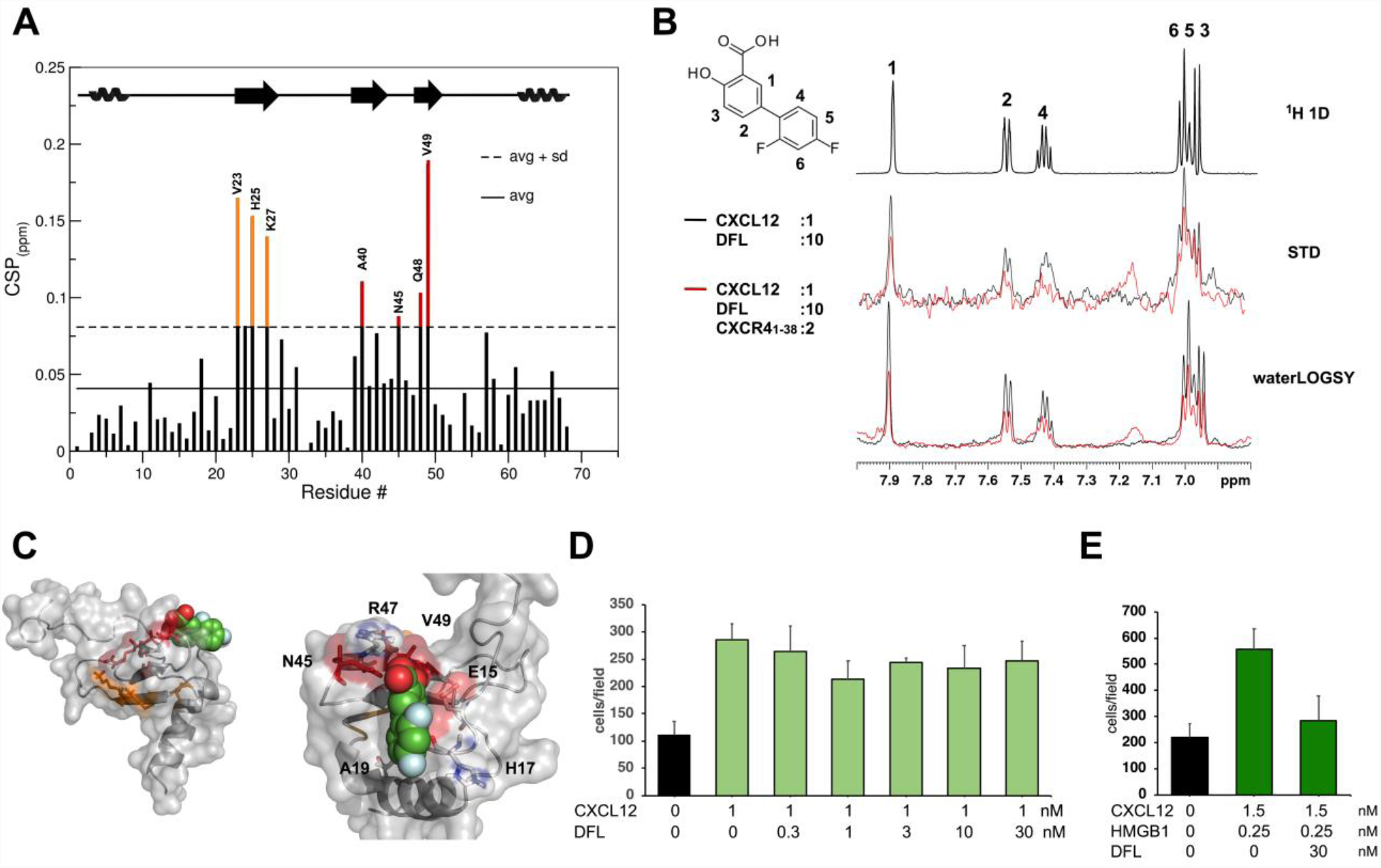
Binding of DFL to CXCL12. A) Histogram showing the CSPs of ^15^N-labelled CXCL12 amides (∼0.1 mM) upon addition of ten-fold excess of DFL. Missing residues are prolines. Elements of secondary structure are depicted on top. B) Top: ^1^H spectrum of DFL, where the numbered peaks correspond to DFL protons (left). Middle: superimposition of STD spectra obtained for 0.5 mM DFL with 0.05 mM CXCL12 (black line) and upon addition of 0.2 mM of CXCR4_1-38sY21_ (red line). Bottom: waterLOGSY spectra obtained for 0.5 mM DFL with 0.05 mM CXCL12 (black line) upon addition of 0.2 mM of CXCR4_1-38sY21_ (red line). C) HADDOCK model of interaction of DFL (CPK representation) with CXCL12 (grey surface and cartoon). CXCL12 residues with CSP > avg + sd, located around the sY21 binding site and on the β1 strand are in red and orange, respectively. Zoom in the binding site, CXCL12 residues (sticks) involved in hydrophobic and electrostatic interactions with DFL (CPK) are explicitly labelled (right). D) DFL does not inhibit CXCL12 induced chemotaxis. 3T3 fibroblasts were subjected to chemotaxis assays in Boyden chambers; 1 nM CXCL12 or no chemoattractant was added in the lower chamber, together with the indicated concentrations of DFL. The data represent the avg ± sd of three replicates, and the migration in the presence or absence of DFL is not significantly different (one-way ANOVA plus post tests). E) DFL inhibits chemotaxis toward the HMGB1/CXCL12 heterocomplex. The data represent the avg ± sd of three replicates. Migration in the presence or absence of DFL is significantly different (p<0.001, one-way ANOVA plus Tukey’s post test).

Accordingly, data driven docking calculations indicated that the carboxylate and the hydroxyl of DFL are respectively well suited for polar interactions with the sidechains of R47 and E15 located within the sY21 binding pocket (Fig. 2C).

Ligands targeting the sY21 binding pocket are also known to allosterically trigger CXCL12 dimerization upon binding (Ziarek *et al*, 2013). Thus, we hypothesized that CSPs involving the β1 strand (Fig. 2A,C) might be due to protein dimerization. Indeed, the increase of CXCL12 correlation time upon DFL addition (τ_c_ of CXCL12_free_ = 5.5 ± 0.15 ns, τ_c_ of CXCL12_DFL_ = 6.3 ± 0.4 ns, Appendix Fig. S7), as assessed by heteronuclear NMR relaxation experiments, supports the notion that DFL promotes CXCL12 self-association.

Cell migration experiments, however, showed that DFL was unable to inhibit CXCL12-induced chemotaxis at concentrations where it inhibited chemotaxis induced by HMGB1 (Fig. 2D). Thus, neither binding to HMGB1 alone or CXCL12 alone can justify the inhibition of chemotaxis. However, DFL might also bind to and/or interfere with the HMGB1/CXCL12 heterocomplex. Indeed, DFL significantly reduced the migration of 3T3 fibroblasts chemotaxis towards the preformed HMGB1/CXCL12 heterocomplex (Fig. 2E). To verify biophysically that DFL interferes with the HMGB1/CXCL12 heterocomplex, we first produced the heterocomplex by titrating unlabeled CXCL12 (up to 0.2 mM) into ^15^N labelled HMGB1 (0.1 mM) (Fig. 3A,B top), and then added DFL (Fig. 3D,E). As already reported (Schiraldi *et al*, 2012), complex formation induced the reduction or disappearance of peak intensities of HMGB1 amide resonances, indicative of micromolar affinities in intermediate exchange regime on the NMR time scale (Fig. 3B). These observations were in agreement with K_d_ ∼ 4 ± 0.4 μM, as measured by MST (Fig. 3C).

**Figure 3.**
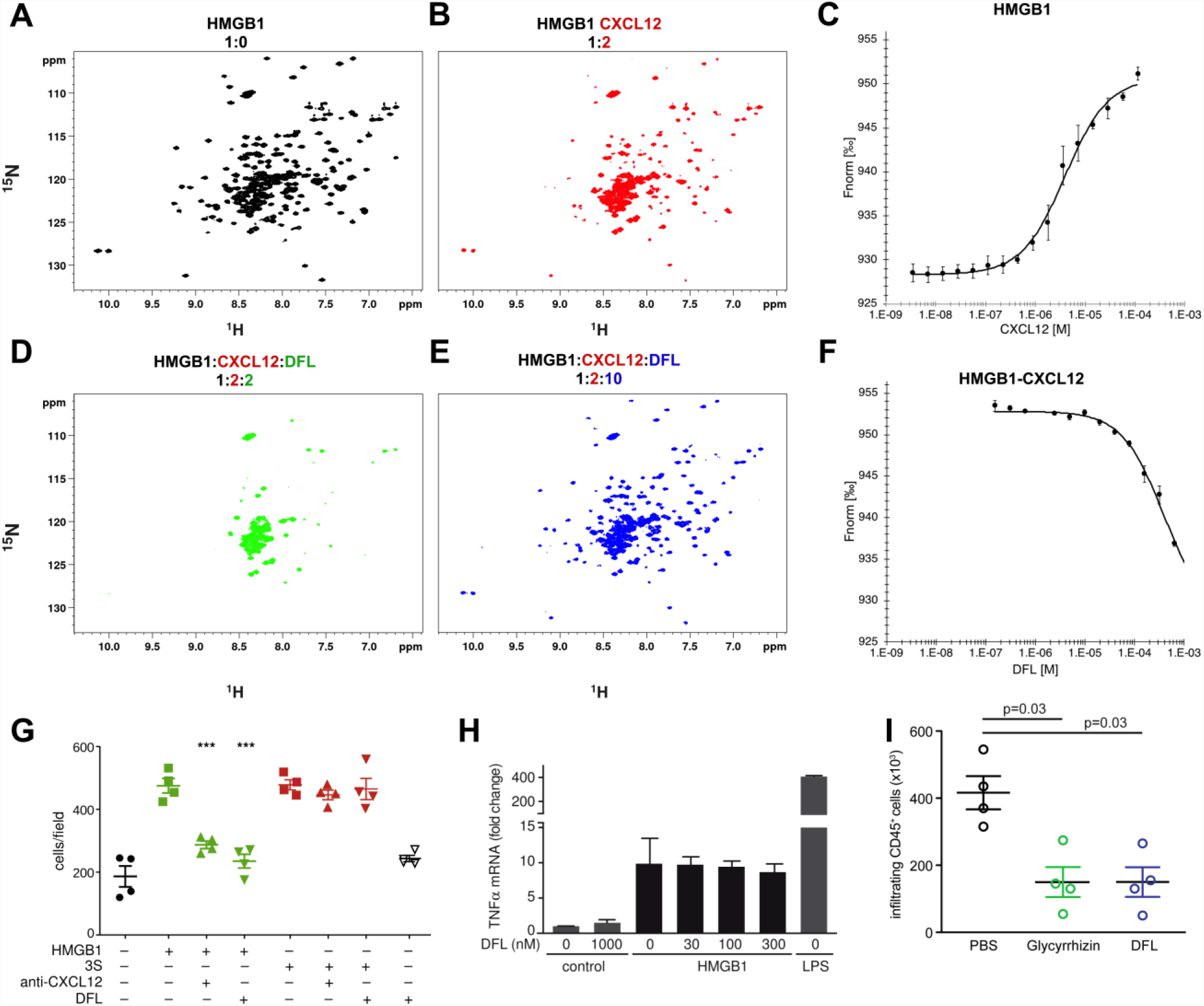
Effect of DFL on HMGB1/CXCL12 heterocomplex. A) ^1^H-^15^N HSQC HMGB1 (0.1 mM) spectrum without (black) and B) with a two-fold excess of CXCL12, C) MST curves of CXCL12 titrated into fluorescently labelled HMGB1; the signal increases from ∼900 (free) to ∼930 a.u. (bound) yielding an apparent K_d_ of 4 ± 0.4 μM. H-^15^N HSQC HMGB1 (0.1 mM) with CXCL12 (0.2 mM) upon addition of D) two-fold and E) ten-fold excess of DFL. F) MST competition experiment: the heterocomplex is preformed using 50 nM and 50 μM of HMGB1 and CXCL12, respectively, and measurements were performed in the presence of increasing concentrations of DFL. Decreasing MST signal (from ∼930 to ∼910 a.u.) upon DFL titration indicates that CXCL12 has been displaced from HMGB1. *n*=3; error bars represent ± sd. G) DFL does not inhibit chemotaxis towards 3S. 3T3 fibroblasts were subjected to chemotaxis assays in Boyden chambers; 1 nM fully reduced HMGB1 (green) or 3S (red), or no chemoattractant (black), were added in the lower chamber, together with 1 µg/ml of anti-CXCL12 monoclonal antibody or 30 nM DFL. The data represent the avg ± sd of four replicates. ***, p<0.001 relative to migration towards HMGB1, one-way ANOVA plus Tukey’s post test; the migrations towards 3S are not significantly different among each other. H) DFL does not affect the cytokine-inducing activities of disulfide HMGB1. Mouse Bone Marrow Derived Macrophages were activated or not for 3 h with 3 µg/ml disulfide HMGB1 (∼100 nM) or 10 ng/ml LPS, in the presence of the indicated concentrations of DFL. The bars represent the mean ± sd (n=3 biological replicates). I) Quantification of CD45+ cells isolated with immunobeads from injured muscles 6 hours after damage. Before damage mice were treated either with PBS (Phosphate buffered saline) vehicle, Glycyrrhizin or DFL. n=4 mice per group, two independent experiments. Bars and error bars represent avg ± sd (statistics: Kruskal-Willis plus post-tests).

Intriguingly, HMGB1 residues involved in interactions with CXCL12 and DFL partially coincide (Appendix Fig. S8), suggesting that DFL might interfere with the surface of interaction of HMGB1 with CXCL12. Upon addition of 0.2 mM DFL to the preformed ^15^N-HMGB1/CXCL12 heterocomplex we observed a drastic line broadening in the ^1^H-^15^N HSQC spectrum, with the disappearance of the majority of HMGB1 peaks (Fig. 3D, green). These line broadening effects were conceivably associated to multiple equilibria involving free HMGB1, HMGB1 bound to DFL, HMGB1 bound to CXCL12 and the ternary complex involving HMGB1, CXCL12 and DFL. The remaining peaks, in the typical random coil region of the C-terminal HMGB1 acidic tail, were not involved in interactions with either CXCL12 or DFL. By ultimately adding a ten-fold excess of DFL the ^1^H-^15^N HSQC spectrum of HMGB1 was restored (Fig. 3E, blue, Appendix Figure S9), indicating that CXCL12 was displaced and the heterocomplex disrupted. Notably, some HMGB1 cross-peaks were shifted with respect to free HMGB1 due to binding of DFL (Appendix Fig. S9). In MST experiments addition of increasing concentrations of DFL to a preformed HMGB1/CXCL12 complex lead to a sharp transition at 380 ± 17 μM, consistent with the disassembly of fluorescently labelled HMGB1 from the heterocomplex (Fig. 3F).

In line with the evidence that DFL affects the HMGB1/CXCL12 heterocomplex, we predicted that DFL would not affect the chemoattractant activity of 3S, a mutant of HMGB1 where all cysteines are replaced by serines. 3S has the same chemoattractant properties as HMGB1, but does not need to form a complex with CXCL12 (Tirone *et al*, 2018). Indeed, both a monoclonal antibody against CXCL12 and DFL inhibited HMGB1-induced chemotaxis, but neither was effective on chemotaxis induced by 3S (Fig. 3G).

HMGB1 containing a disulfide bond between cysteines 22 and 44 induces cytokine production by binding to receptor TLR4 (He *et al*, 2018). Conversely, disulfide-containing HMGB1 cannot form a complex with CXCL12 (Schiraldi *et al*, 2012). Thus, if DFL specifically targets the HMGB1/CXCL12 heterocomplex, it should not interfere with the transcription and release of pro-inflammatory cytokines specifically triggered by disulfide HMGB1. Indeed, addition of DFL did not affect the transcription of TNFα in mouse macrophages stimulated with disulfide HMGB1 (Fig. 3H); likewise, DFL did not affect cytokine induction by disulfide HMGB1 in human macrophages (Appendix Fig. S10). This is at variance with the effect of Salicylic Acid, an active metabolite of aspirin, which inhibits the cytokine-inducing activities of disulfide HMGB1 (Choi *et al*, 2015).

Next, we tested *in vivo* the ability of DFL to inhibit the migration of inflammatory cells into injured muscle, which also depends on the HMGB1/CXCL12 heterocomplex (Tirone *et al*, 2018). Muscle injury was induced by a single dose of cardiotoxin in the tibialis anterior muscle of C57Bl/6 wild type mice, and the infiltration of leukocytes (CD45+ cells) in injured muscle was assessed 6 hours later (Schiraldi *et al*, 2012). DFL significantly decreased the recruitment of CD45+ cells into the injured muscle compared to vehicle-treated mice (Fig. 3I). As previously reported (Schiraldi *et al*, 2012), Glycyrrihizin also inhibited recruitment of leukocytes into injured muscle (Fig. 3I); we thus tested whether its inhibitory activity was mechanistically similar to that of DFL. Indeed, NMR titration experiments clearly showed that Glycyrrhizin binds directly to CXCL12, affecting a binding surface similar to that targeted by DFL (Appendix Fig. S11). Most importantly, Glycyrrhizin was able to disrupt the HMGB1-CXCL12 complex, as shown in ^15^N-HSQC experiments by the recovery of ^15^N HMGB1 resonances upon addition to a preformed heterocomplex (Appendix Fig. S12). Of note, in strong analogy to DFL, Glycyrrhizin is not an inhibitor of CXCL12 induced chemotaxis (Schiraldi *et al*, 2012).

Taken together, these data support a multi-step molecular mechanism whereby DFL and Glycyrrhizin inhibit the chemotactic activity of HMGB1 by targeting the HMGB1/CXCL12 heterocomplex. The unexpected combinatorial binding activity of HMGB1 inhibitors might in part reconcile the unusual differences observed between their inhibition activity and the affinities to their single targets. Herein, DFL appears to work as a multi-target drug (Bolognesi & Cavalli, 2016; Anighoro *et al*, 2014), displaying a low affinity towards its multiple targets but resulting in high efficiency through their simultaneous engagement (Korcsmáros *et al*, 2007).

Salicylates and their derivatives are potent and widely used anti-inflammatory drugs with a wide spectrum of modulating activities on cyclooxygenases (Warner *et al*, 1999), NF-kB (Kopp & Ghosh, 1994), AMPK (Hawley *et al*, 2012) and CBP/p300 acetyltransferase activity (Shirakawa *et al*, 2016). Results presented in this work add to the variety of mechanisms in which DFL is involved. Importantly, DFL has been and is widely employed, and the understanding of its mode of action can inform and improve its clinical usage. Notably, the use of DFL should be discouraged during the healing of damaged tissue, as we showed that the HMGB1/CXCL12 heterocomplex favors muscle, bone or liver regeneration after damage (Tirone *et al*, 2018).

In conclusion, the ability of DFL to selectively interfere with the HMGB1/CXCL12 inflammatory axis through the simultaneous binding to both HMGB1 and CXCL12 offers unprecedented structural/functional insights into the anti-inflammatory activity of DFL as a NSAID. These insights also show that protein-protein interactions within the HMGB1-CXCL12 heterocomplex are druggable with high specificity and selectivity, thus opening new strategies for the rational design of novel inhibitors of this crucial inflammatory axis.

## Materials and Methods

Recombinant HMGB1 constructs, including Box A (residues 1–89), Box B (residues 90– 175), full-length HMGB1 (residues 1-214) were produced by expressing constructs in a pETM-11 vector (EMBL, Heidelberg, DE). Proteins contained an N-terminal 6His-tag, removable by cleavage with TEV protease. After expression and cleavage with TEV protease, the proteins have a residual N-terminal three-residue tag (GAM). Box A and Box B proteins were expressed in *E. coli* BL21 (DE3) cells, whereas HMGB1 was expressed in pLys BL21 (DE3) *E. coli* cells. Cells were grown at 37°C until the optical density at 600 nm reached 0.8 absorbance units. Gene expression was induced by the addition of isopropyl β-D-1-thiogalactopyranoside (IPTG) to a final concentration of 1 mM. After 18 hours of incubation at 25°C with shaking, cells were harvested by centrifugation. The cells were re-suspended in lysis buffer (20 mM Tris-HCl pH 8.0, 150 mM NaCl, 10 mM imidazole, 2 mM β-mercaptoethanol, 0.2% NP-40, Complete EDTA-free protease inhibitor (Roche) and lysed by sonication. Cell debris was removed by centrifugation at 11,000 rcf for 45 min at 4°C. The soluble 6His-tagged proteins were purified from the supernatant by affinity chromatography using Ni^2+^-NTA agarose resin (Qiagen, Hilden, Germany). After several washing steps, proteins were eluted in 20 mM Tris pH 8.0, 150 mM NaCl, 300 mM imidazole, 2 mM β-mercaptoethanol. The 6His-tag was removed by overnight incubation at 4°C with TEV protease. During incubation, the sample was dialyzed against 20 mM Tris pH 8.0, 150 mM NaCl, 2 mM β-mercaptoethanol for Box A and Box B, whereas for HMGB1 the dialysis buffer was 20 mM Tris pH 8.0, 20 mM NaCl, 2 mM β-mercaptoethanol. Uncleaved 6His-tagged protein and TEV protease were then removed by repassing the sample over Ni^2+^-NTA resin. HMGB1 was further purified on a HitrapQ ion-exchange column using buffer A (20 mM Tris pH 8, 20 mM NaCl, 2 mM β-mercaptoethanol) and buffer B (20 mM Tris pH 8, 1 M NaCl, 2 mM β-mercaptoethanol) to create a linear gradient of NaCl. HMGB1 elutes at 500 mM NaCl. Box A and B samples were purified by gel filtration on a Superdex-75 column (Amersham Biosciences, Milan, Italy) equilibrated in 20 mM phosphate buffer (pH 7.3), 150 mM NaCl, 1 mM dithiothreitol (DTT). For MST experiments, the 6His-tag was not removed. Uniformly ^15^N-and ^15^N/^13^C-labelled proteins were prepared using M9 minimal bacterial growth media appropriately supplemented with ^15^N-labelled ammonium chloride and ^13^C-labelled glucose. Unlabeled and uniformly ^15^N-and ^15^N/^13^C-labelled CXCL12 were provided by HMGBiotech (Milan, Italy). HMGB1 R23AK27A mutant was produced as described in (Choi *et al*, 2015). For biophysical measurements, all HMGB1 constructs, after expression and purification, were dialyzed against NMR buffer, containing 20 mM phosphate buffer pH 7.3, 150 mM NaCl, 1 mM DTT. CXCL12 was dialyzed against a buffer containing 20 mM phosphate buffer pH 6, 20 mM NaCl.

Proteins concentrations were determined measuring the absorbance at 280 nm considering molar extinction coefficients of 9970, 10810, 21430 and 8700 M^-1^ cm^-1^ for Box A, Box B, HMGB1 and CXCL12, respectively.

Proteins used for cell-based and *in vivo* assays were provided by HMGBiotech (Milan).

Diflunisal (5-(2,4-difluorophenyl)-2-hydroxybenzoic acid) was purchased from Sigma-Aldrich.

### Peptide synthesis and purification

The CXCR4_1-38sY21_ peptide with sequence MEGIDIYTSDNYTEEMGSGDY(Sulfo)DSMKEPAFREENANFNK was synthesized using the Fmoc method (Fields & Noble, 1990) and purified by preparative reversed-phase chromatography (RP-HPLC). Details on peptide synthesis and purification are reported in Appendix.

### NMR Measurements

NMR spectra were recorded at 298 K on a Bruker Avance 600 MHz spectrometer (Karlsruhe, Germany) equipped with a triple-resonance TCI cryoprobe with an x, y, z-shielded pulsed-field gradient coil. Spectra were processed with Topspin^TM^ 3.2 (Bruker) and analyzed with CcpNmr Analysis 2.3. (Vranken *et al*, 2005) ^1^H-^15^N-HSQC assignments of HMGB1 and its constructs were taken from the BMRB databank (accession numbers: 15148, 15149). The ^1^H-^15^N-HSQC assignment of CXCL12 was obtained from the BMRB databank (accession number 16143) (Veldkamp *et al*, 2009) and confirmed *via* acquisition of 3D HNCA, CBCA(CO)NH experiments. Sidechain assignment of Box A (Box B) in the presence of DFL (stoichiometric ratio 1:2) was obtained through the analysis of 3D HNCA, CBCA(CO)NH, CBCANH, H(CCO)NH, CC(CO)NH and HCCH-TOCSY experiments and 2D ^1^H-^1^H TOCSY (mixing time: 60 ms) and NOESY (mixing time: 120 ms) spectra. Intermolecular nuclear Overhauser effect (nOes) between DFL and Box A (Box B) were obtained from 3D ^13^C-NOESY-HSQC with no evolution on ^13^C dimension (2048 × 1 × 256 increments) experiments with ^15^N/^13^C filter in F1 (mixing time 200 ms); protein and ligand concentration were 0.8 mM and 1.6 mM, respectively, in D_2_O.

#### Titrations

For NMR titrations, at each titration point a 2D water-flip-back ^1^H-^15^N-edited HSQC spectrum was acquired with 2048 (160) complex points for ^1^H (^15^N), respectively, apodized by 90° shifted squared (sine) window functions and zero filled to 256 points for indirect dimension. Assignment of the labelled proteins in the presence of the ligands (DFL or unlabeled protein) was obtained following individual cross-peaks through the titration series. For each residue the weighted average of the ^1^H and ^15^N chemical shift perturbation (CSP) was calculated as CSP = [(Δδ^2^HN + Δδ^2^N/25)/2]^1^/^2^ (Grzesiek *et al*, 1996). In titrations of ^15^N HMGB1 with unlabeled CXCL12, because of extensive line broadening, the effect of the binding was monitored measuring the variation of ^1^H-^15^N peak intensities ratios I/I_o_ upon CXCL12 addition, where I_o_ and I are peak intensities in free and bound HMGB1, respectively. Intensities ratios I/I_o_ values were normalized on the highest value.

DFL titrations have been performed on ^15^N HMGB1 constructs (full length protein, Box A, Box B, 20 mM phosphate buffer, pH 7.3, 20 mM NaCl, 1 mM DTT) and on ^15^N CXCL12 (20 mM phosphate buffer, pH 6, 20 mM NaCl) adding 0.5, 1, 2, 3, 5, 10 equivalents of DFL to the labelled proteins. In order to minimize dilution and NMR signal loss, titrations were carried out by adding small aliquots of concentrated DFL (10 mM in 20 mM phosphate buffer, pH 7.3, 150 mM NaCl) to the ^15^N labelled protein samples (0.1 mM).

CXCL12 (1 mM stock solution, 20 mM phosphate buffer, pH 6, 20 mM NaCl) was titrated adding 0.5, 1, 1.5, 2 equivalents into 0.1 mM ^15^N-labelled HMGB1 (20 mM phosphate buffer, pH 6, 20 mM NaCl, 1 mM DTT). DTT is required to maintain HMGB1 in its fully reduced form. We suspected that DTT might break important disulfide bridges within CXCL12; however, control ^1^H-^15^N HSQC spectra acquired in the presence of 1 mM DTT indicated that ^15^N-labelled CXCL12 in these conditions was stable for at least 4 hours, a time scale which was well within the time required for NMR titrations.

NMR-based antagonist-induced dissociation assays (Krajewski *et al*, 2007) were performed titrating DFL on the ^15^N-HMGB1/CXCL12 heterocomplex (ratio 1:2) adding 1, 2, 3, 5, 10 equivalents of ligand to the heterocomplex. Similar NMR-based antagonist induced dissociation assays were performed titrating Glycyrrhizin into the ^15^N-HMGB1/CXCL12 heterocomplex (ratio 1:2) adding 0.25, 0.5, 0.75, 1, 1.5, 2.5 and 4 equivalents of ligand to the heterocomplex.

Glycyrrhizin (Acros Organics) titration on ^15^N CXCL12 (20 mM phosphate buffer, pH 6, 20 mM NaCl) has been performed adding 0.25, 0.5, 1, 2 equivalents of ligand to the labelled proteins using a 10 mM stock solution (20 mM phosphate buffer, pH 7.3, 150 mM NaCl). *Saturation Transfer Difference and Water-ligand observed via gradient spectroscopy.* STD and waterLOGSY experiments have been performed on 0.5 mM DFL in the presence of 0.05 mM HMGB1 or CXCL12 in NMR buffer. STD experiments were acquired using a pulse scheme (Bruker pulse sequence: stddiffesgp.3) with excitation sculpting with gradients for water suppression and spin-lock field to suppress protein signals. The spectra were acquired using 128 scans, a spectral width of 9600 Hz, 64K data points for acquisition. For protein saturation, a train of 60 Gaussian shaped pulses of 50 ms was applied, for a total saturation time of 3 s. Relaxation delay was set to 3 s. On-and off-resonance irradiations were set at 0 ppm and at 107 ppm, respectively. STD spectra were obtained by internal subtraction of the on-resonance spectrum from the off-resonance spectrum.

WaterLOGSY experiments were acquired using a pulse scheme as described (Dalvit *et al*, 2000) with excitation sculpting and flip-back for water suppression. The spectra were acquired using 128 scans, 32K data points for acquisition, mixing time was set to 1 s.

Competition experiments have been performed comparing STD and waterLOGSY performed on 0.5 mM DFL with 0.05 mM CXCL12 and upon addition of 0.2 mM of CXCR4_1-38sY21_ peptide.

#### Dissociation constant estimation

The apparent dissociation constant between DFL and ^15^N-CXCL12 was estimated from least-squares fitting of CSPs (9 residues) as a function of total DFL concentration according to the equation:

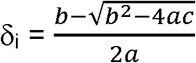

with a=(K_a_/δ_b_) [P_t_], b= 1+K_a_([L_ti_]+[P_t_]), and c=δ_b_K_a_[L_ti_], where δi is the absolute change in chemical shift for each titration point, [L_ti_] is the total DFL concentration at each titration point, [P_t_] is the total protein concentration, K_a_ =1/K_d_ is the association constant, and δ_b_ is the chemical shift of the resonance in the complex. K_d_ and δ_b_ were used as fitting parameters using the Xmgrace program (http://plasma-gate.weizmann.ac.il/Grace/).

#### Relaxation Experiments

Relaxation experiments were performed on ^15^N labelled CXCL12 (0.1 mM in 20 mM phosphate buffer, 20 mM NaCl, pH 6) in the absence and in the presence of 10-fold excess DFL at 298 K.

Heteronuclear {^1^H}^15^N nuclear Overhauser enhancement, longitudinal and transversal ^15^N relaxation rates (R_1_, R_2_) were measured using standard 2D methods (Farrow *et al*, 1994). For R_1_ and R_2_ experiments a duty-cycle heating compensation was used(Yip & Zuiderweg, 2005); the two decay curves were sampled at 12 (ranging from 50 to 2000 ms) and 11 (from 12 to 244 ms) different time points, respectively. Both R_1_ and R_2_ experiments were collected in random order and using a recovery delay of 2.5 s. R_1_ and R_2_ values were fitted to a 2-parameter exponential decay from the intensities using the fitting routine implemented in the CcpNmr program; duplicate measurements were used to allow a statistical analysis of the uncertainties (Skelton *et al*, 1993). HSQC spectra measured in an interleaved fashion with and without 4 s of proton saturation during recovery delay were recorded for the {^1^H}^15^N heteronuclear nOe experiments. The corresponding values were obtained from the ratio between saturated and unsaturated peaks intensities. The uncertainty was estimated through the ratio between the standard deviation of the noise in both saturated and unsaturated spectra divided by the intensity of the respective peaks. The correlation time (τ_c_) was estimated from R_2_/R_1_ according to the following equation (Bloembergen *et al*, 1948)

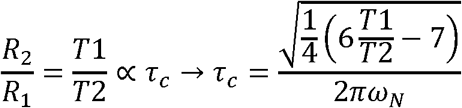

Only residues corresponding to non-overlapping peaks, with heteronuclear nOe > 0.65 were used, residues with anomalous R_2_ were excluded (Barbato *et al*, 1992).

### Docking models

Molecular docking of DFL on Box A (residues G3-Y77), Box B (A93-G173) (coordinates were extracted from the PDB structure with code: 2YRQ) and CXCL12 (K1-K68, coordinates extracted from the PDB structure with code: 4UAI) were performed using the data-driven software HADDOCK 2.2 (Dominguez *et al*, 2003; van Zundert *et al*, 2016) following the classical three-stage procedure which includes: (1) randomization of orientations and rigid body minimization, (2) simulated annealing in torsion angle space, and (3) refinement in Cartesian space with explicit water. Ambiguous interaction restraints (AIRs) were defined as follows: residues with CSP > avg + sd or displaying intramolecular nOes were used to define active residues, whose solvent accessible surface neighbors were set as passive (Appendix Table S1). In the case of CXCL12, only the residues located around the sY21 binding site (violet CSP in Fig. 2D) were set as active (Appendix Table S1), as STD competition experiments of DFL in the presence of CXCR4_1-38sY21_ demonstrated that they both compete for the sY21 binding site. In the case of Box A (Box B), intermolecular nOes were included as unambiguous restraints in the calculations only in the semi-flexible refinement stage, setting the maximum distance of the nOe H pairs to 5 Å (Appendix Table S1).

Optimized parameters for liquid simulation (OPLS) were used for the protein (protein-allhdg5-4 and protein-allhdg5-4-caro). The geometric coordinates and parameters for DFL were calculated and optimized using the PRODRG server (Schüttelkopf & van Aalten, 2004). Calculations generated 1000, 1000, 500 structures for the rigid body docking (it0), the semi-flexible refinement (it1) and the explicit solvent refinement (water), respectively. The final 500 structures obtained after water refinement were scored according to their HADDOCK score. The latter (defined as HADDOCKscore = 1.0 E_vdW_ + 0.2 E_elec_ + 1.0 E_desolv_ + 0.1 E_AIR_) is a weighted combination of van der Waals (vdW) and electrostatic energy terms (Lennard–Jones and Coulomb potentials), empirical desolvation term (Fernández-Recio *et al*, 2004) and ambiguous interaction restraint energy term, which reflects the accordance of the model to the input restraints.

HADDOCK models were clustered (Daura *et al*, 1999) based on their interface root mean square deviation (rmsd), setting the cutoff and the minimum number of models in a cluster to 1.8 Å and 10 for the boxes, and 2.5 Å and 10 for CXCL12, respectively. Proteins were aligned and fitted on the backbone of active residues reported in Table S1. The rmsd of DFL was calculated only on the heavy atoms of the entire scaffold.

To remove any bias of the cluster size on the cluster statistics, the final overall score of each cluster was calculated on the four lowest HADDOCK scores models in that cluster. For each protein the cluster with the best fitting with respect to the experimentally-driven restraints (lowest number of violations) and the best HADDOCK score (cluster 1 for Box A, cluster 5 for Box B and CXCL12) was selected (Appendix Fig. S13, S14 and S15).

The analysis of the docking calculations was performed applying in-house python and tcl scripts. Molecular images were generated by PyMOL Molecular Graphics System, Version 2.0 Schrödinger, LLC.

### MST experiments

MST experiments were performed at 24°C on a NanoTemper® Monolith NT.115 instrument with red filters, using 40% LED power and 60% MST power. All the experiments were carried out using 6His-tagged HMGB1, non-covalently labelled with the NT647 fluorescence dye (Bartoschik *et al*, 2018). MST buffer contained 20 mM phosphate buffer pH 7.3, 150 mM NaCl, 0.05% Tween, 1 mM DTT.

For binding assays, 16-points titration series of DFL or CXCL12 were prepared, whereby the ligand dilutions were generated as a 1:2 dilution of the stock solution using MST buffer; a constant amount of labelled HMGB1 (50 nM) was added to all dilution steps. Maximum concentrations of DFL and CXCL12 in the titrations series were 5 mM and 217 μM, respectively. Complex samples were incubated for 15 minutes before loading into NanoTemper premium capillaries.

Competition experiments were carried out pre-forming a complex between labelled 6His-tagged HMGB1 (50 nM) and unlabeled CXCL12 (10 μM, *i.e.* 2 times the estimated K_d_) (Jerabek-Willemsen *et al*, 2011). For 16-points titration series of DFL, serial 1:2 dilutions of the DFL stock solution were made into MST buffer, and a constant amount of pre-formed heterocomplex was added to all dilution steps. All samples were incubated for 15 minutes before measurements. Maximum concentration of DFL in the titrations series was 0.63 mM. Addition of DFL induced the recovery of the MST signal of HMGB1 towards the unbound state value (Fig. 3C). Attempts to reach saturation failed, as higher DFL concentrations (up to 5 mM) induced aggregation phenomena, as assessed by the bumpiness of the MST traces (data not shown).

For all MST experiments data points were the average of three measurements (error bars correspond to standard deviation). Data analyses were carried out using NanoTemper analysis software using the K_d_ model fitting for the binding assays and Hill model for competition experiments.

### Migration experiments

For fibroblast chemotaxis, modified Boyden chambers were used with filters (pore diameter 8 µm; Neuro Probe) coated with 50 µg/mL fibronectin (Roche). 50,000 cells in 200 µL were added to the upper chamber. Serum-free DMEM as negative control, HMGB1 and/or other molecules were added to the lower chamber at the indicated concentration, and then cells were left to migrate for 3 hours at 37°C. Cells were fixed with ethanol and stained with Giemsa Stain (Sigma), then non-migrating cells were removed with a cotton swab. All assays were done at least in triplicate. The migrated cells were acquired with Zeiss Imager M.2 microscope at 10x magnification, then evaluated with an automated counting program.

### Assay for Cytokine Induction on mouse BMDMs

Bone Marrow Derived Macrophages (BMDMs) were obtained as described (Zhang *et al*, 2008) and stimulated as described in the legend to Fig. 3. Total RNAs were isolated using the Illustra RNAspin Mini kit (GE Healthcare), and complementary DNAs (cDNAs) were obtained by reverse transcription with oligo(dT) primers (Invitrogen, Carlsbad, CA, USA) and SuperScript II Reverse Transcriptase (Invitrogen) following the manufacturers’ instructions. Quantitative real-time PCR was performed using a LightCycler480 (Roche Molecular Diagnostics), in triplicates, using SYBR Green I master mix. The ΔCt method was used for quantification, and the β-actin gene was used for normalization.

The sequence of the primers was

TNFα forward: CTTCTCATTCCTGCTTGTGG

TNFα reverse: GCAGAGAGGAGGTTGACTTTC

Beta-actin forward: AGACGGGGTCACCCACACTGTGCCCATCTA

Beta-actin reverse: CTAGAAGCACTTGCGGTGCACGATGGAGGG

### Human macrophages

Peripheral blood mononuclear cells (PBMCs) were isolated from buffy coats of donor blood (Hospital of Magenta, Italy) by Ficoll gradient centrifugation (Lymphoprep, AXIS-SHIELD). CD14+ monocytes were isolated by positive immunoselection (CD14 MicroBeads, Miltenyi Biotec, Germany) according to the manufacturer’s instructions, and differentiated into macrophages using X-Vivo medium supplemented with 1% heat inactivated human serum, GM-CSF and M-CSF.

#### *In vivo* injury model

To assess the initial leukocyte recruitment after muscle injury, mice were injected intravenously with 200 µg of DFL (Sigma Aldrich) or Glycyrrhizin (Acros Organics) 3 hours before muscle injection with 50 µL of 15 µM cardiotoxin (Latoxan). After 6 hours, the injured Tibialis Anterior muscles were collected and dissociated in RPMI 1640 containing 0.2% collagenase B (Roche Diagnostics) at 37°C for 1 hour. CD45^+^ cells were purified by magnetic cell sorting by using anti-CD45 beads (Miltenyi Biotec) according to the manufacturer’s instructions and quantified by Countess (Invitrogen).

## Acknowledgements

This work was supported by an iCARE Fellowship funded by AIRC and Marie Curie Actions – People – COFUND (Project code 16258), and by AIRC (IG18623; IG21440). We gratefully acknowledge Dr. Francesca Viganò and Dr. Katarzyna Walkiewicz (NanoTemper Technologies GmbH, Munich), Dr. Delia Tarantino (University of Milan) for technical assistance and discussion in MST experiments and Dr. Samuel Zambrano for useful discussions.

## Author contribution

F.D.L. performed the NMR experiments, computational studies, analyzed the data and prepared the manuscript. G.Q. performed the NMR experiments, M.T. performed the in vivo experiments and analyzed the data. V.M., A.P. and M.C. expressed and purified recombinant proteins. F.D.M. performed the cell migration experiments. A. G. synthesized and purified the peptide. R.M. performed the cytokine induction experiments. G.M. and M.E.B. directed the study and were involved in all aspects of the experimental design, data analysis and manuscript preparation. All authors critically reviewed the text and figures.

## The Paper Explained

### PROBLEM

HMGB1 is a key molecule that both triggers and sustains inflammation following infection or injury, and is involved in a large number of inflammatory and autoimmune diseases, and in certain types of cancer. Thus, it is appealing to develop drugs that target HMGB1 specifically. Moreover, HMGB1 has two broad activities: inflammatory cell recruitment, and induction of chemokines and cytokines. Targeting one class of activities and not the other one is even more attractive.

### RESULTS

In this work we show that Diflunisal (DFL), an aspirin-like nonsteroidal anti-inflammatory drug that has been extensively used for many decades under the commercial name of Dolobid, inhibits HMGB1 *in vitro* and *in vivo* and blocks the recruitment of inflammatory cells at low nanomolar concentrations. Importantly, DFL does not interfere with chemokine/cytokines induction.

We have also worked out the unusual mode of action of DFL. Recruitment of inflammatory cells is known to depend on the formation of a heterocomplex between HMGB1 and the chemokine CXCL12, which in turn activates CXCR4, a G-protein coupled receptor (GPCR). Using a variety of techniques, we show that DFL binds directly to *both* HMGB1 and CXCL12, and targets and disrupts the HMGB1-CXCL12 heterocomplex.

### IMPACT

We have thus at least partially clarified the mode of action of DFL. Notably, since we find that DFL targets the HMGB1-CXCL12 heterocomplex, we suggest that it should not be given to patients recovering from injury or trauma. Most importantly, our work shows that protein-protein interactions within the HMGB1-CXCL12 heterocomplex are druggable, thus opening new strategies for the rational design of novel inhibitors.

## Appendix.

The Appendix contains supplementary methods for peptide synthesis and Supplementary Figures and Tables.

Correspondence and requests for materials should be addressed to G.M and M.E.B.

