## Supplementary Information for "Diflunisal targets the HMGB1/CXCL12 heterocomplex and blocks immune cell recruitment"

### Appendix

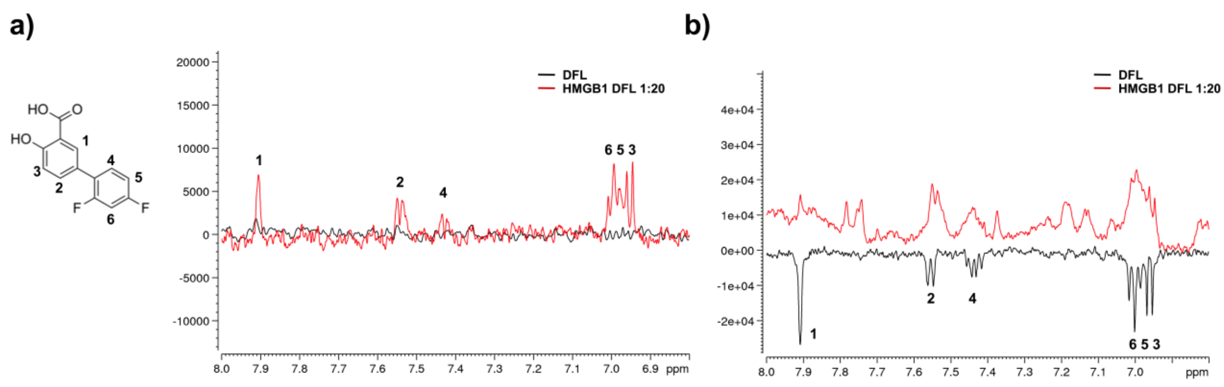


**Appendix Figure S1.** a) Saturation Transfer Difference (STD) results obtained for 1 mM DFL alone (black line) and with 0.05 mM HMGB1 (red line) in 20 mM phosphate buffer, 150 mM NaCl, 1 mM DTT pH 7.3 (on resonance: 0 ppm; saturation time: 3 s). STD signals with different intensities are observed for all protons in the presence of HMGB1, indicating that DFL directly binds to HMGB1. The numbered peaks correspond to proton resonance assignments. b) WaterLOGSY spectra obtained for 1 mM DFL alone (black line) and in complex with 0.05 mM HMGB1 (red line). All DFL protons display signals inversion, indicating binding to HMGB1.

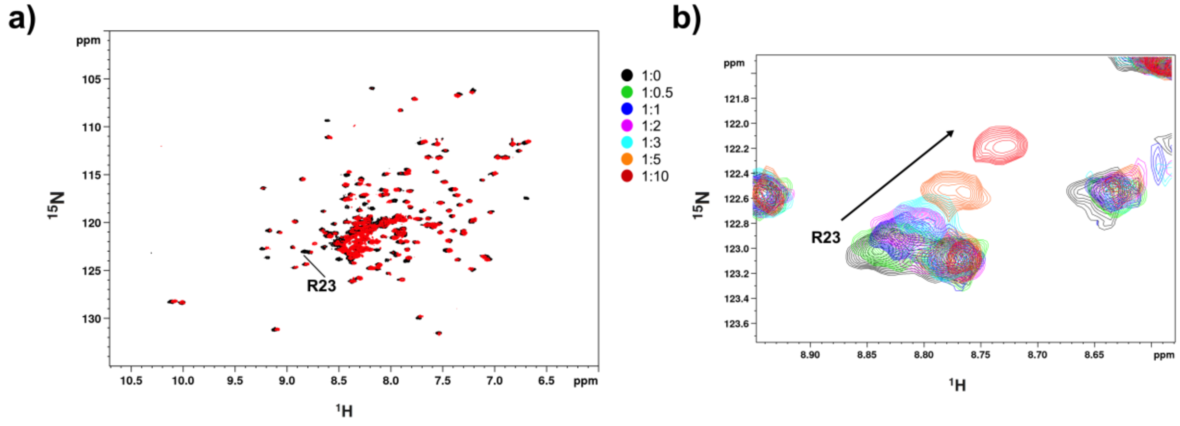


**Appendix Figure S2.** a) Superimposition of the $^{1}$H-$^{15}$N HSQC spectra of HMGB1 (~0.1mM, pH 7.3, phosphate buffer, 1 mM DTT) without (black) and with ten-fold excess of DFL (red). b) Selected region of the superimposition of $^{1}$H-$^{15}$N HSQC spectra of HMGB1 during the titration with DFL (0.5, 1, 2, 3, 5 and 10 equivalents) showing the displacement of the peak associated to R23 along the titration.

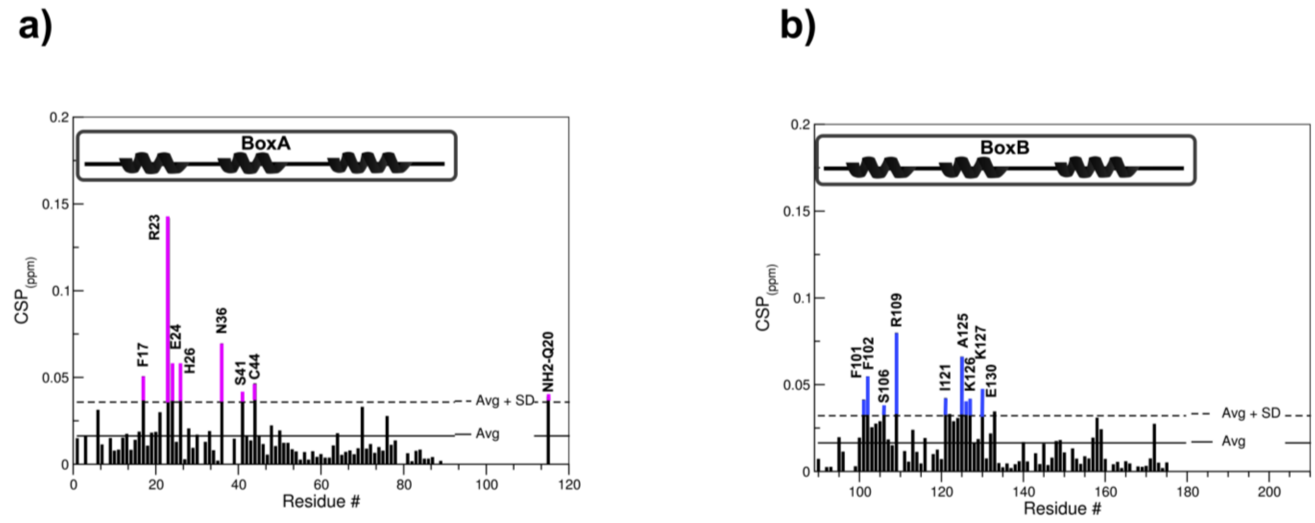


**Appendix Figure S3.** Histograms of CSPs (ppm) of (a) Box A) and (b) Box B amides upon addition of ten-fold excess of DFL. CSPs > avg + sd are highlighted in magenta (Box A) and in blue (Box B). α-helices are schematically represented on top.

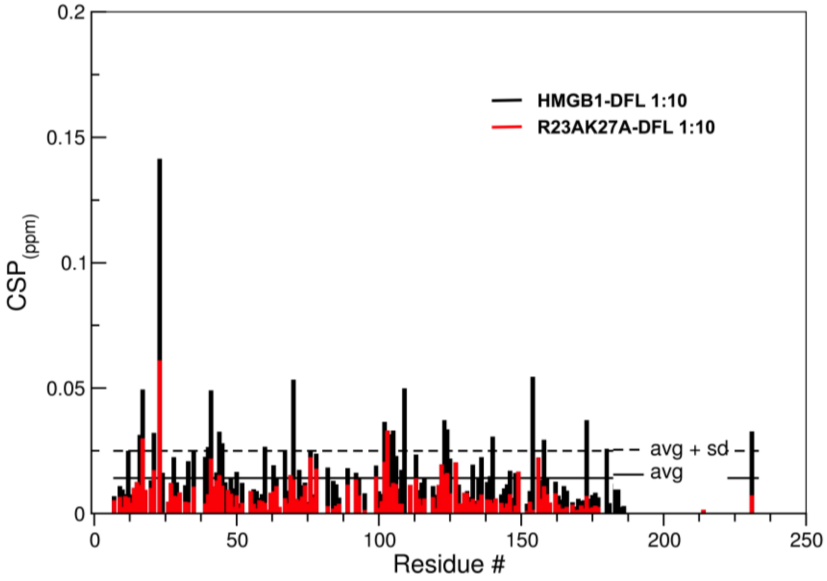


**Appendix Figure S4.** Histograms of CSPs (ppm) of HMGB1 (black) and R23AK27A mutant (red) amides upon addition of ten-fold excess of DFL.

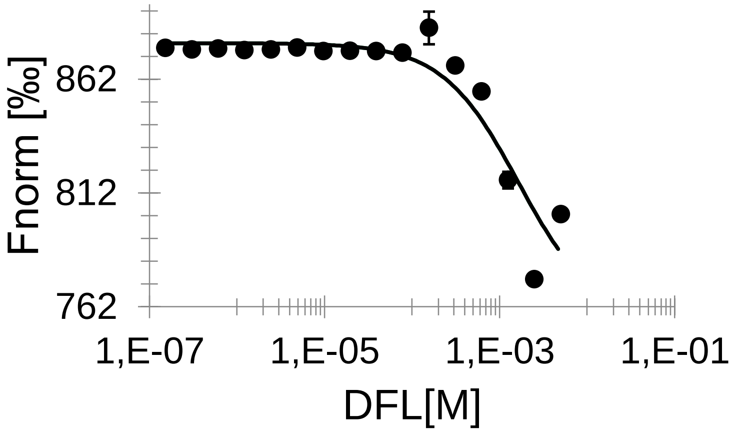


**Appendix Figure S5.** Dose-response curve of HMGB1-DFL interaction followed by MST. DFL concentrations ranged from 5 mM to 153 nM on 50 nM labelled 6His-HMGB1. The $K_d$ is 1.6 ± 0.8 mM (n=3, error bars correspond to sd). Saturation of the curve could not be reached because of solubility issues for DFL stock solutions > 10 mM. (Fnorm=normalized fluorescence).

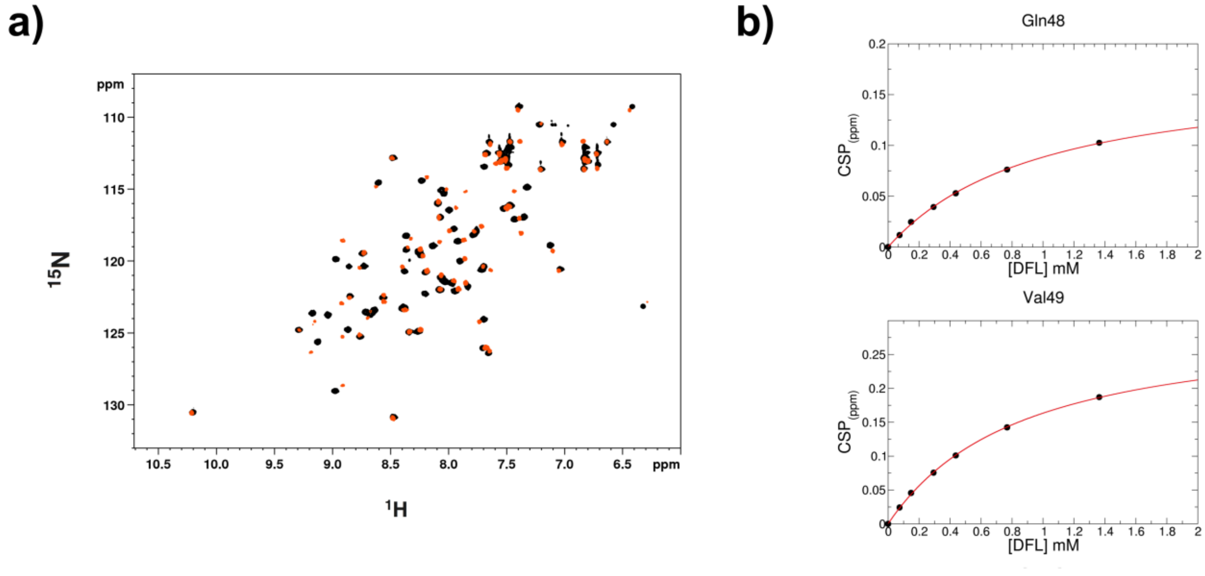


**Appendix Figure S6.** a) Superposition of $^{1}$H-$^{15}$N HSQC spectra of $^{15}$N CXCL12 (~0.1 mM, pH 6, phosphate buffer) without (black) and with ten-fold excess of DFL (orange). b) Weighted average of Q48 and V49 amide $^{1}$H and $^{15}$N chemical-shift changes in the presence of increasing concentration of DFL. Nonlinear curve fitting yields $K_d$ = 802 ± 102 μM.

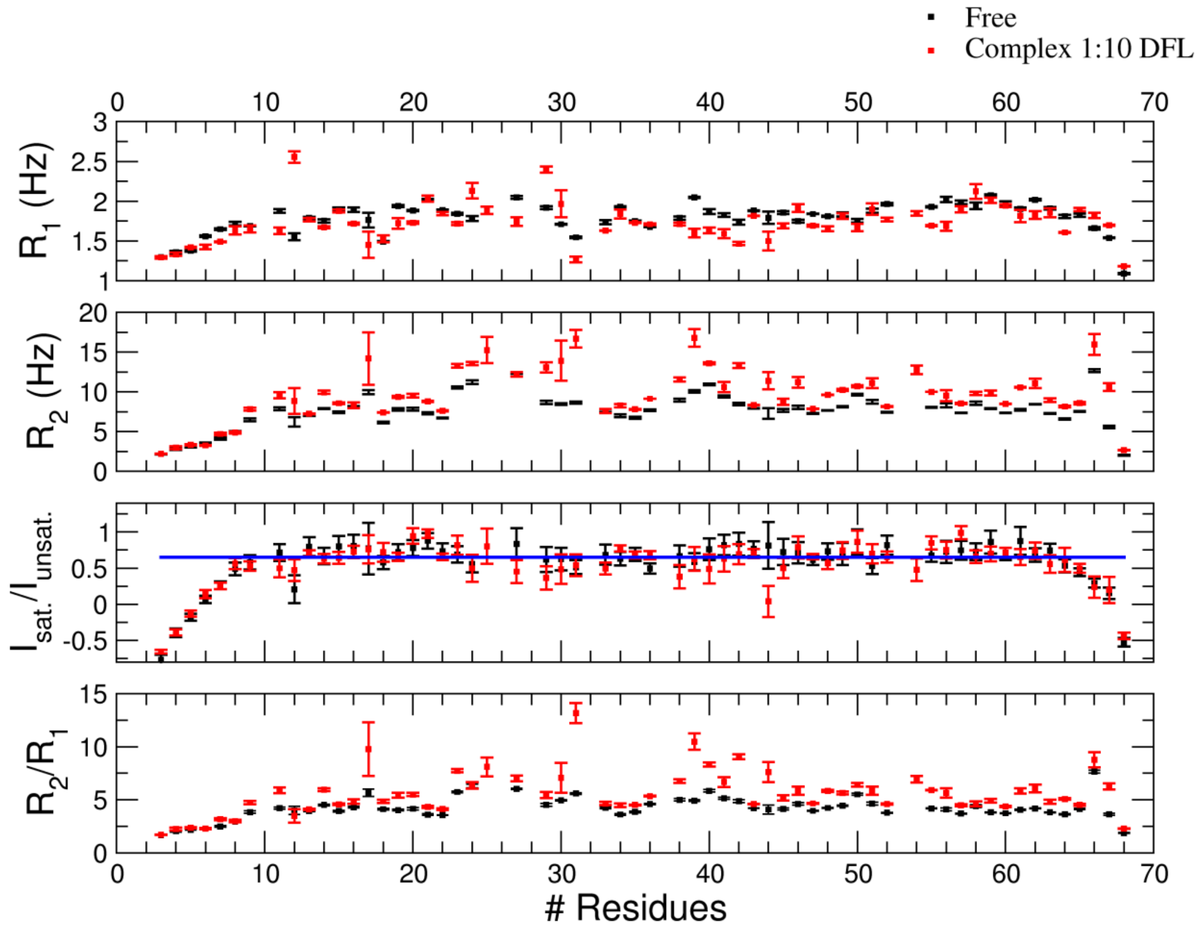


**Appendix Figure S7.** $^{15}N$ $R_1$, $R_2$ relaxation rates and $\{^1H\}$–$^{15}N$ heteronuclear nOe values of free CXCL12 (0.1 mM, black) and in complex with 10-fold excess of DFL (red). The blue horizontal line indicates the ratio of saturated over unsaturated intensity ($I_{sat}/I_{unsatur}$) with a threshold value of 0.65. Measurements were performed at 298 K.

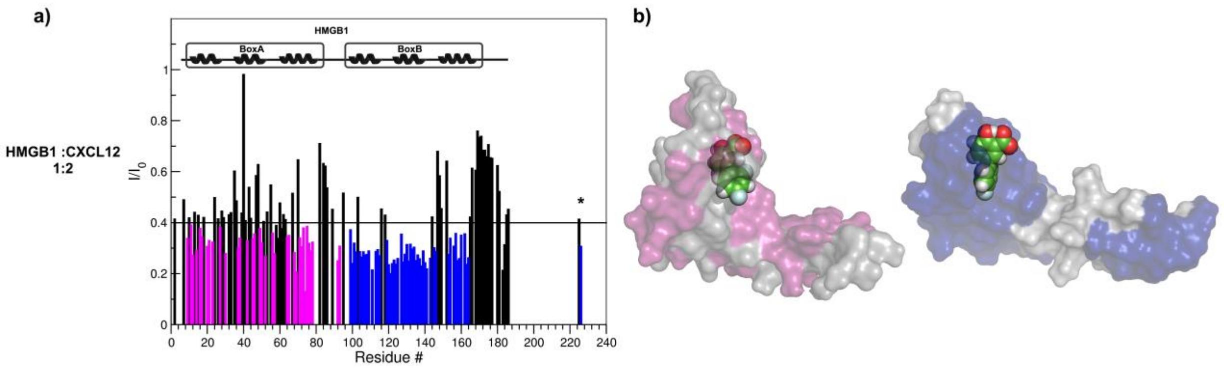


**Appendix Figure S8.** a) Histograms showing the $I/I_0$ peak intensity ratio of HMGB1 amides (~0.1 mM, pH 6, phosphate buffer, 1 mM DTT) with and without two-fold excess of CXCL12 (bottom). $I_o$ and I are peak intensities in free and bound HMGB1, respectively. Residues showing significant decrease of $I/I_0$ ratio ($I/I_0$ ratio < avg) upon CXCL12 binding are represented with magenta (Box A) and blue (Box B) histograms. $\alpha$-helices are schematically represented on the top of the histogram. b) Surface representations of Box A-DFL and Box B-DFL molecular models are reported where the residues showing significant decrease of $I/I_0$ ratio ($I/I_0$ ratio < avg) upon CXCL12 binding are mapped in magenta and blue, respectively.

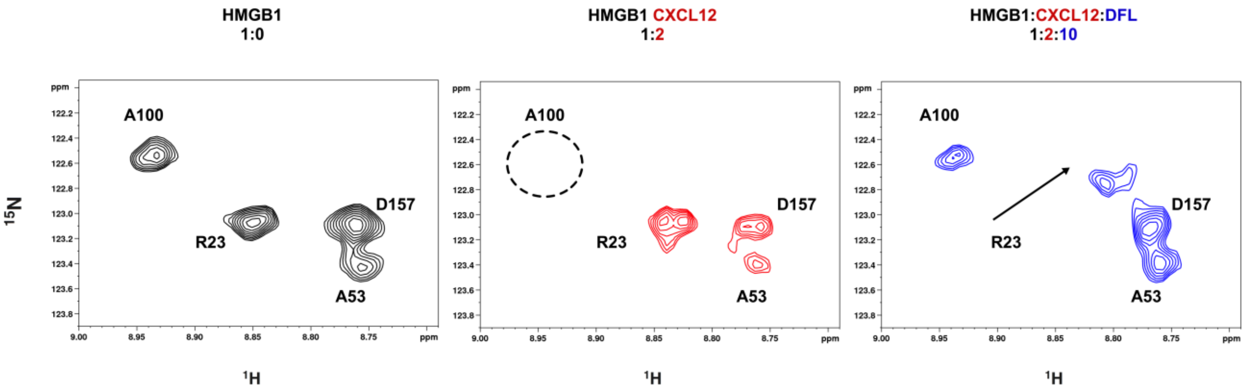


**Appendix Figure S9.** Selected region of $^1H$-$^{15}N$-HSQC spectra of $^{15}N$ HMGB1 (~0.1 mM, pH 6, phosphate buffer, 1 mM DTT) without (black), with two-fold excess of CXCL12 (red) and ten-fold excess of DFL (blue). In the presence of CXCL12 the HMGB1 spectrum undergoes line broadening and some peaks (*e.g.* A100 in Box B, black dotted circle) disappear, indicating complex formation. The A100 peak reappears in the $^1H$-$^{15}N$-HSQC spectrum of $^{15}N$ HMGB1 upon addition of DFL, whereas the sites involved in binding to DFL are shifted (*e.g.* R23).

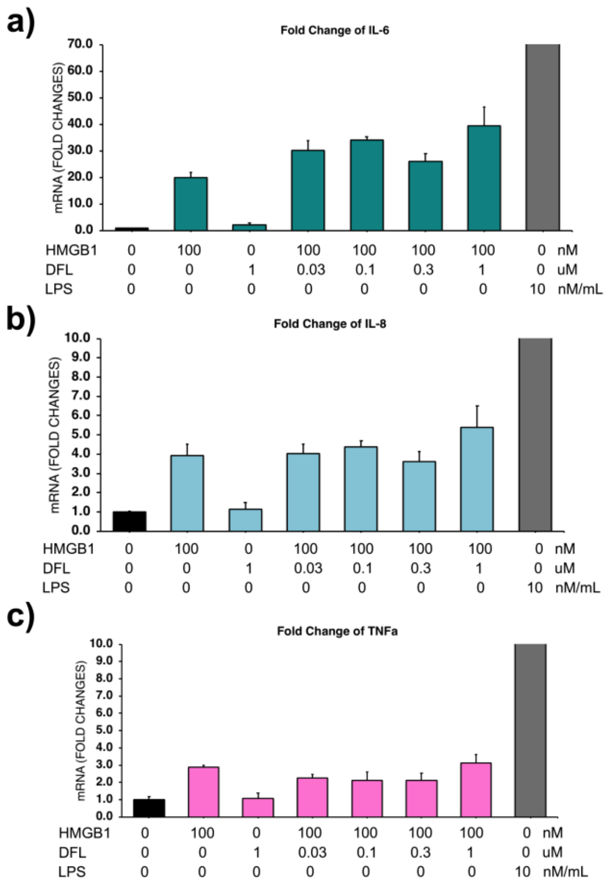


**Appendix Figure S10.** DFL does not affect the cytokine-inducing activities of disulfide HMGB1 on human macrophages. Human macrophages were obtained as described (Venereau *et al*, 2012) and activated or not for 3 h with 3 µg/ml disulfide HMGB1 (~100 nM) or 10 ng/ml LPS, in the presence of the indicated concentrations of DFL. The bars represent the mean ± sd (n=3 biological replicates). DFL does not produce a statistically difference in cytokine/chemokine transcription following exposure to disulfide HMGB1 (one-way ANOVA plus post tests). Total RNAs were isolated using the Illustra RNAspin Mini kit (GE Healthcare), and complementary DNA (cDNA) was obtained by retro-transcription with Oligo(dT) primers (Invitrogen, Carlsbad, CA, USA) and SuperScript II Reverse Transcriptase (Invitrogen) following the manufacturers' instructions. Quantitative real-time PCR was then performed using LightCycler480 (Roche Molecular Diagnostics), in duplicates, using SYBR Green I master mix and the following primers:

β-actin: 5'-TGACGGGGTCACCCACACTGTGCCC-3', and 5'-CTAGAAGCATTGCGGTGGAC GATGG-3';

TNF-α: 5'-AGCCCATGTTGTAGCAAACC-3' and 5'-AGGACCTGGGAGTAGATGAGG-3';

IL-6: 5’-TACCCCCAGGAGAAGATTCC-3’ and 5’-TTTTCACCAGGCAAGTCTCC-3;

IL-8: 5’-TGCCAAGGAGTGCTAAAG-3’ and 5’-CTCCACAACCCTCTGCAC-3’.

The ΔCt method was used for quantification, and the β-actin gene was used for normalization.

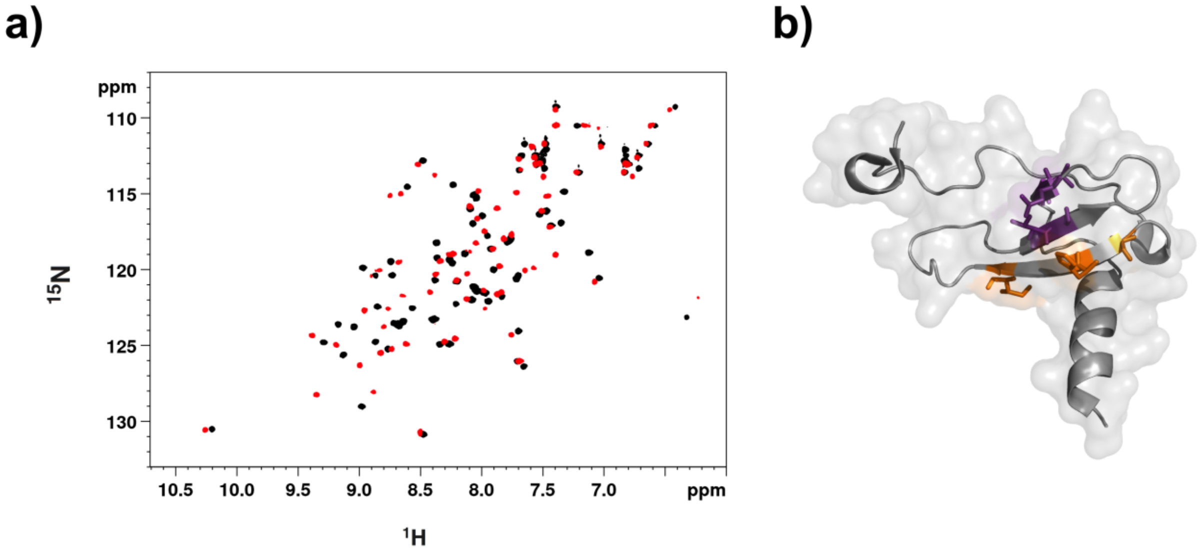


**Appendix Figure S11.** a) Superimposition of the $^{1}H$-$^{15}N$ HSQC spectra of $^{15}N$ CXCL12 (~0.1mM, pH 6, phosphate buffer) without (black) and with 2-fold excess of Glycyrrhizin (red). b) Mapping of CXCL12 residues with CSP > avg + sd, located around the sY21 binding site and on the β1 strand are in violet and orange, respectively.

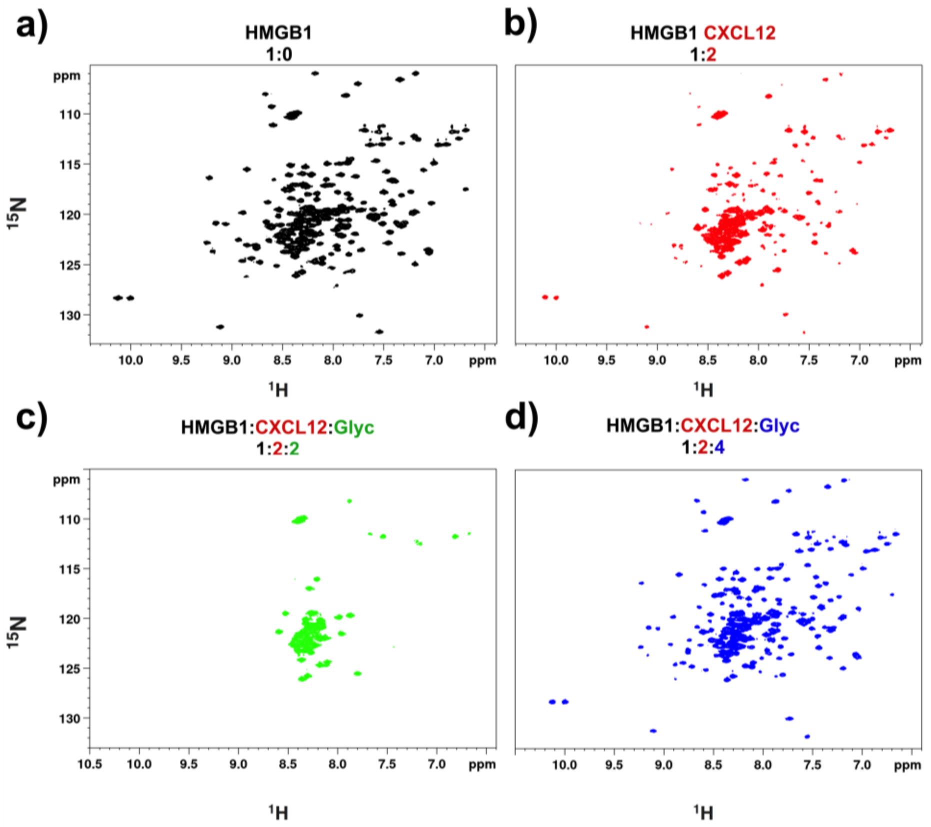


**Appendix Figure S12.** a) $^{1}H$-$^{15}N$ HSQC of $^{15}N$ HMGB1 (~0.1 mM) spectrum without (black), b) with a two-fold excess of CXCL12 (red), c) upon addition of two-fold (green) and d) four-fold excess of Glycyrrhizin (Glyc) (blue).

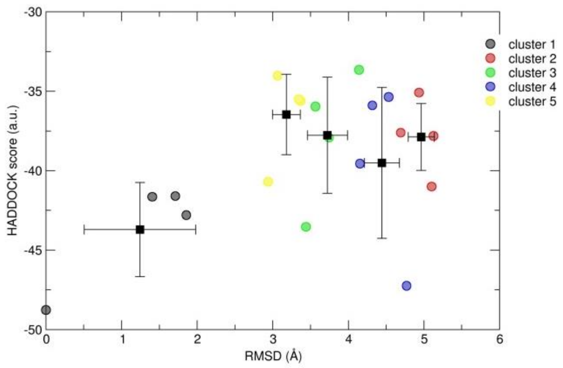


**Supplementary Figure 13.** HADDOCK score *versus* rmsd from the lowest Haddock energy complex structure between Box A and DFL. Circles correspond to the four best structures in each cluster, the cluster averages with the standard deviation are indicated with the black squares and bars.

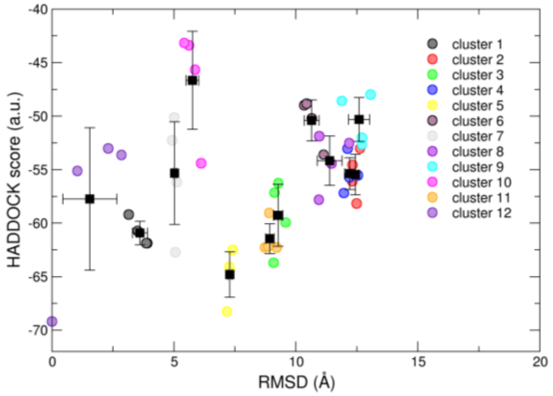


**Appendix Figure S14.** HADDOCK score *versus* rmsd from the lowest HADDOCK energy complex structure between Box B and DFL. Circles correspond to the four best structures in each cluster, the cluster averages with the standard deviation are indicated with the black squares and bars.

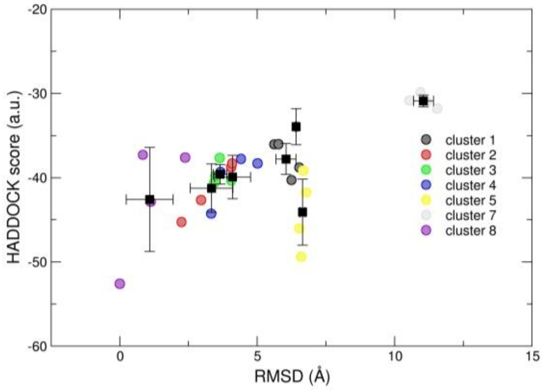


**Appendix Figure S15.** HADDOCK score *versus* rmsd from the lowest HADDOCK energy complex structure between CXCL12 and DFL. Circles correspond to the four best structures in each cluster, the cluster averages with the standard deviation are indicated with the black squares and bars.

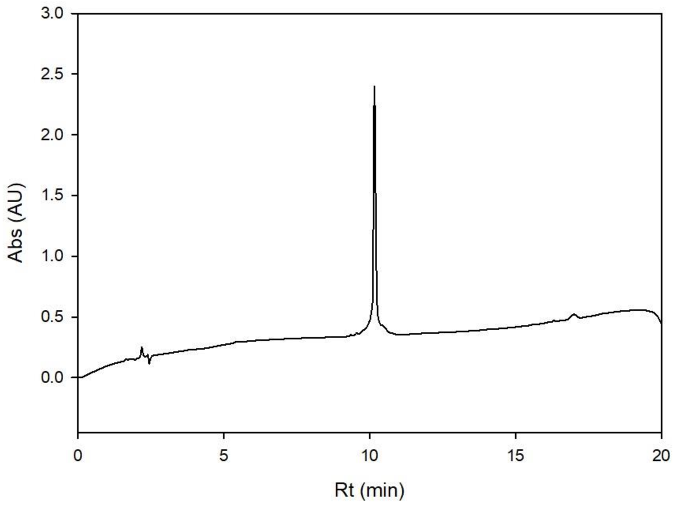


**Appendix Figure S16.** RP-HPLC trace of sulfopeptide CXCR4$_{1\text{-}38sY21}$. Analytical gradient: 70%A to 0% over 14 min.

**Table S1.** Residues used as AIRs, Unambiguous (nOe) restraints used in HADDOCK calculations.

| | Ambiguous | | Unambiguous[a,b] |
|---|---|---|---|
| Domain | Active | Passive | |
| Box A | A16, F17, Q20, T21, R23, H26, V35, P40, S41, C44, S45 | M12, S13, S14, T15, A16, F37, W48 | HE1$_{H26}$-H1$_{DFL}$; HD*$_{V35}$-H3 $_{DFL}$ or HE*$_{V35}$-H3 $_{DFL}$; HZ*$_{P40}$-H1/2$_{DFL}$, or HD*$_{P40}$-H1/2$_{DFL}$ or HE*$_{P40}$-H1/2$_{DFL}$ |
| Box B | C105, R109, I112, K113, I121, V124, A125, K127, L128, W132, K140 | F101, F104, S106 G118, L119 S120, D123, K126, L128, G129 | HD*$_{I112}$-H1$_{DFL}$; HG*$_{V124}$-H6/12 $_{DFL}$; HD*$_{L128}$-H5 $_{DFL}$ |
| CXCL12 | A40, L42, N45, Q48, V49 | E15, H17, V18, A19, N22, N44, R47 | *n.d.* |

*n.d.* Not determined

[a] OPLS *force field* nomenclature of protein hydrogens

[b] DFL hydrogens (Fig. 1 and Appendix Fig.S1)

### Supplementary methods

#### Peptide synthesis and purification

The $CXCR4_{1\text{-}38sY21}$ sulfo-peptide with sequence MEGIDIYTSDNYTEEMGSGDY(Sulfo)DSMKEPAFREENANFNK was assembled by stepwise microwave-assisted Fmoc-SPPS on a Biotage ALSTRA Initiator+ peptide synthesizer, operating in a 0.12 mM scale on a HMPB-CM resin (0.5 mM/g). Resin was swelled prior to use with an NMP/DCM mixture. Activation and coupling of Fmoc-protected amino acids were performed using Oxyma 0.5 M / DIC 0.5 M (1:1:1), with a 5-equivalent excess over the initial resin loading. 1 mL of 1 M LiCl was added to the reaction mixture and coupling steps were performed for 30 minutes at 50°C. Fmoc-Tyr($SO_3$-Np)-OH was coupled at room temperature for 4 hours. Deprotection steps were performed by treatment with a 20% piperidine solution in DMF at room temperature (1 x 10 min, 40°C). Following each coupling or deprotection step, peptidyl-resin was washed with DMF (4 x 4 mL). Upon complete chain assembly, the peptide was cleaved from the resin using a TFA 92.5%, water 2.5%, thioanisole 2.5%, TIS 2.5% mixture (2 hours, RT). Following precipitation in cold diethyl ether, crude peptide was collected by centrifugation and washed with further cold diethyl ether to remove scavengers. Peptide was then dissolved in 50% aqueous acetonitrile 0.07% TFA buffer and purified by preparative RP-HPLC.

*RP-HPLC analysis and purification*

Analytical and semi-preparative reversed phase high performance liquid chromatography (RP-HPLC) were carried out on a Shimadzu Prominence HPLC system equipped with a diode array detector for analytical purposes. Preparative HPLC was performed using a Shimadzu Prominence preparative system (UV detector). A Shimadzu Shim-pack GWS 5µ C18 90 Å column (150 x 4.6 mm) was used for analytical runs and a Shimadzu Shim-pack G15 10µ C18 90Å (250 x 20 mm) for peptide purification. Data were recorded and processed with a Labsolutions software. HPLC eluent A = $H_2O$/ 3 % $CH_3CN$ / 0.07 % TFA; HPLC eluent B = 70 % $CH_3CN$/ 30 % $H_2O$/ 0.07 % TFA. Peptides purification was achieved by preparative RP-HPLC at a flow rate of 14 mL/min using a 100% A to 30% B gradient over 40 min. Pure RP-HPLC fractions (Appendix Figure S16) (>95%) were combined and lyophilized.

*Sulfotyrosine neopentyl protection removal*

Tyr(SO3-np) peptide was dissolved (5 mg/mL) in 50 mM $Na_2HPO_4$ and pH adjusted to 7.2. The peptide solution was incubated overnight to allow sulfotyrosine neopentyl group removal (HPLC monitoring), and the resulting product HPLC-purified (Appendix Fig. S16).

*Electro-spray ionization mass spectrometry (ESI-MS)*

ESI-MS was performed using a Bruker Esquire 3000+ instrument equipped with an electro-spray ionization source and a quadrupole ion trap detector (QITD). Samples were dissolved at a concentration of 0.1 mg/mL in 0.1% formic acid (aq) and injected.
